# Non-invasive ultrasound quantification of Scar Tissue Volume predicts functional changes during tendon healing

**DOI:** 10.1101/460790

**Authors:** Jessica E. Ackerman, Valentina Studentsova, Alayna E. Loiselle

**Author notes:** Corresponding Author: Alayna E. Loiselle, PhD, Center for Musculoskeletal Research, University of Rochester Medical Center, 601 Elmwood Ave, Box 665, Rochester, NY.

## Abstract

Tendon injuries are very common and disrupt the transmission of forces from muscle to bone, leading to impaired function and quality of life. Successful restoration of tendon function after injury is a challenging clinical problem due to the pathological, scar-mediated manner in which tendons heal. Currently, there are no standard treatments to modulate scar tissue formation and improve tendon healing. A major limitation to the identification of therapeutic candidates has been the reliance on terminal end-point metrics of healing in pre-clinical studies, which require a large number of animals and result in destruction of the tissue. To address this limitation, we have identified quantification of Scar Tissue Volume (STV) from ultrasound imaging as a longitudinal, non-invasive metric of tendon healing. STV was strongly correlated with established endpoint metrics of gliding function including Gliding Resistance (GR) and Metatarsophalangeal (MTP) Flexion Angle. However, no associations were observed between STV and tensile mechanical properties. To define the sensitivity of STV to identify differences between functionally discrete tendon healing phenotypes, we utilized S100a4 haploinsufficient mice (S100a4^GFP/+^), which heal with improved gliding function relative to wildtype (WT) littermates. A significant decrease in STV was observed in S100a4^GFP/+^repairs, relative to WT at day 14. Taken together, these data suggest US quantification of STV as a means to facilitate the rapid screening of biological and pharmacological interventions to improve tendon healing, and identify promising therapeutic targets, in an efficient, cost-effective manner.

## Introduction

Tendon injuries disrupt the transmission of forces from muscle to bone, leading to chronic pain, disability and a large socioeconomic burden^1^. Tendon injuries are very common, as there are over 300,000 tendon repair procedures a year in the United States^2^, which result from either acute trauma or chronic tendinopathy. While satisfactory outcome rates vary between tendons, successful restoration of tendon function after injury remains a challenging clinical problem, with up to 40% of flexor tendon injuries healing with functional limitations^3^. Unsatisfactory outcomes of surgical tendon repair procedures are due to the pathological, scar-mediated manner in which tendons heal. Rather than regenerating native tendon tissue, tendons heal via bridging scar tissue composed of a disorganized collagen extracellular matrix (ECM), resulting in mechanical properties that are inferior to native tendon, and increasing the risk of re-injury or rupture. Additionally, scar tissue impairs tendon range of motion (ROM), a particularly problematic complication in the flexor tendons of the hands^3^.

Despite attempts using a variety of biological and tissue engineering approaches, there is currently no consensus therapy to improve outcomes after tendon injury. An insufficient understanding of the underlying mechanisms that contribute to scar-mediated tendon healing is a major impediment to the development of successful therapies. To address this limitation, we developed a murine model of tendon injury and repair that recapitulates many clinical aspects of healing including abundant scar tissue formation and impaired restoration of mechanical properties ^4-6^. However, progress in this field is limited by the absence of cost-effective longitudinal outcome measures of tendon healing. Currently, we quantify impairments in gliding function using end-point analyses of Gliding Resistance and measurement of the Metatarsophalangeal (MTP) flexion angle after loading of the proximal FDL with small weights^4,7^, consistent with large animal^8^ and cadaveric studies^9^. This powerful technique has allowed us to define the temporal course of scar formation in the murine model^4^, and demonstrated the effects of multiple genetic and pharmacological perturbations on the healing process^10-12^. However, these studies require many animals to properly power the study and to get sufficient temporal resolution over the course of scar formation and healing. Thus, current approaches are expensive, time-consuming and do not allow longitudinal evaluation or concomitant assessment of function and tissue morphology in the same specimen. Therefore, our objective was to establish quantification of Scar Tissue Volume from ultrasound (US) images as a longitudinal, non-invasive metric of tendon healing. US-based quantification of tendon excursion will dramatically reduce the number of animals needed by longitudinally assessing a single cohort of animals over the entire course of healing. Furthermore, US-based characterization will provide more flexibility as we characterize novel genetic models and interventions as we can image at many more time-points. In addition, while GR and MTP Flexion allow us to make assumptions about tissue morphology, concomitant assessment of function and morphology are not possible in a single specimen. In contrast, 3D reconstruction and segmentation of US images allow direct assessment and quantification of tissue morphology. Finally, US is an ideal modality to longitudinally assess tendon healing, as it is non-ionizing, and can be easily scaled between pre-clinical and clinical applications.

## Methods

*Animal Ethics:* This study was carried out in strict accordance with the recommendations in the Guide for the Care and Use of Laboratory Animals of the National Institutes of Health. All animal procedures were approved by the University Committee on Animal Research (UCAR) at the University of Rochester (UCAR Number: 2014-004).

*Acute tendon injury and repair*: The following strains of mice were obtained from Jackson laboratories (Bar Harbor, ME): C57BL6/J (#664) and S100a4^GFP/+^and Wildtype (WT) littermates (#012904; B6.129S6-*S100a4^tm1Egn^*/YunkJ). Only female C57Bl/6J mice were used, while male and female S100a4^GFP/+^and WT used in equal proportions across genotypes. At 10-12 weeks of age, mice underwent surgical transection and repair of the flexor digitorum longus (FDL) tendon in the hind paw as we have previously described ^4,10-14^. Briefly, the distal FDL tendon was exposed and transected; two horizontal 8-0 sutures were placed in the intact tendon ends and the tendon was sutured to approximate an end-to-end repair. The tendon was also transected proximally along the tibia at the myotendinous junction to decrease strain on the repair.

*Ultrasound quantification of Scar Tissue Volume*: A high-frequency ultrasound platform (Vevo^®^ 3100, FUJIFILM VisualSonics Inc., Toronto, Canada), with a 70-MHz transducer probe (MX700; FUJIFILM VisualSonics Inc., Toronto, Canada) was used for *in vivo* imaging of the healing tendon. For imaging studies, mice were anesthetized with isoflurane, and placed in the prone position. The right hind paw was gently secured proximal to tibiotalar (ankle) joint with surgical tape. After aligning the ultrasound probe with the tendon at the mid-point of the hind paw, the hind paw was covered in ultrasound gel (Aquasonic 100, Parker, Fairfield, NJ) and imaged. A total of 105 40Fm-thick transverse B mode images were collected across the 4mm region of interest (ROI), which included the entire width of the hind paw. All system settings, including gain (96%), monitor dynamic range (70 dB), and depth (2 cm), were kept constant.

The same cohort of mice (n=9) underwent ultrasound imaging at 7, 14, 20, and 28 days post-surgery, followed by assessment of gliding function and mechanical properties at day 28 (Figure 1A). Two mice were excluded from analysis at day 28 due to failure during testing. An additional cohort of animals (n=7) underwent ultrasound imaging at day 14, followed by sacrifice and assessment of gliding function and mechanical properties.

**Figure 1.**
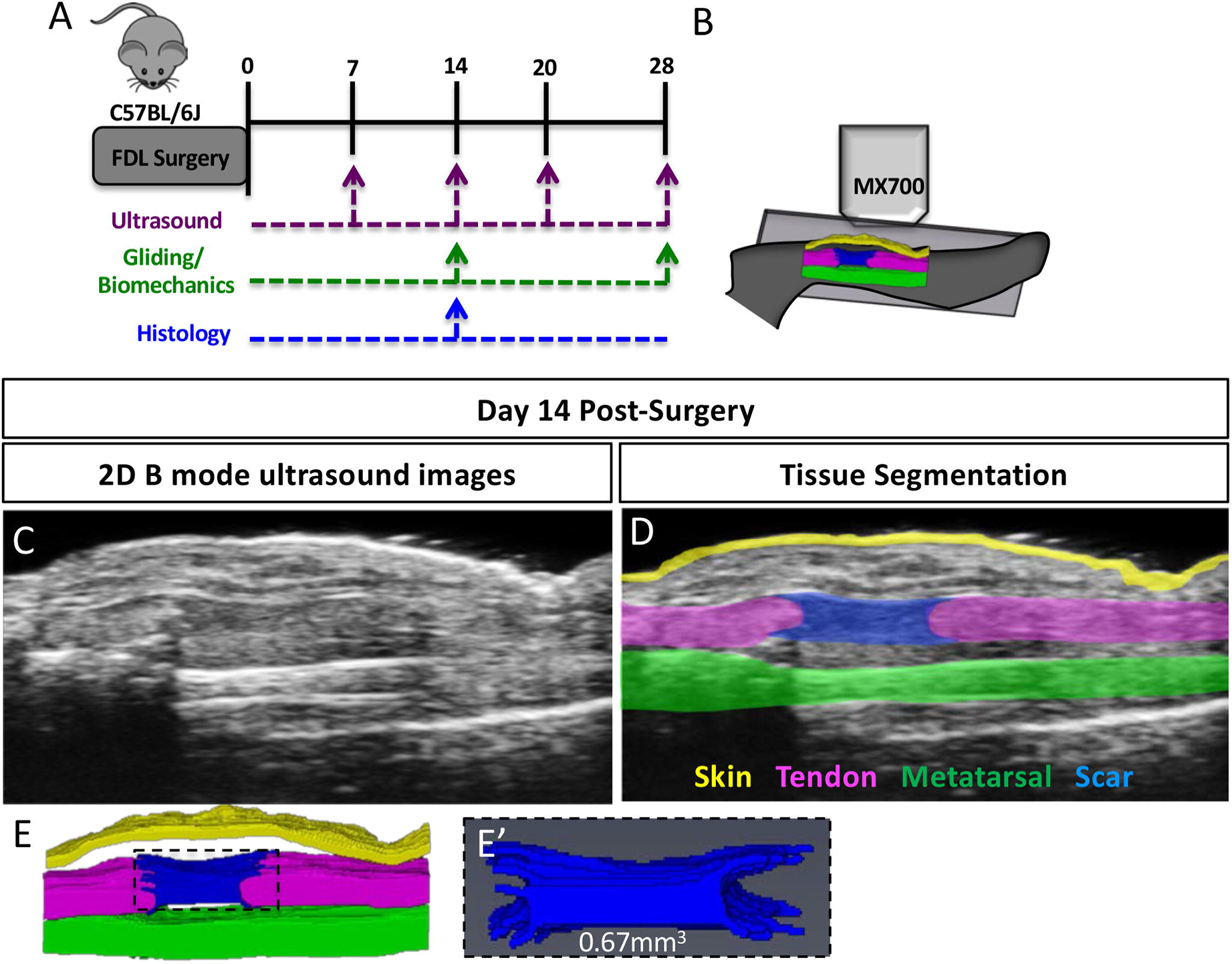
Quantification of Scar Tissue Volume from ultrasound images. (A) Experimental design for longitudinal ultrasound assessment of tendon healing at 7, 14, 20 and 28 days post-surgery, followed by assessment of gliding function at day 28. An additional cohort of animals underwent ultrasound imaging only at day 14, followed by either histological analysis, or assessment of gliding function. (B) Schematic of ultrasound setup showing transverse plane of imaging. (C) 2D B Mode ultrasound image of a healing tendon at day 14 post-surgery. (D) Segmentation of skin (yellow), metatarsal (green), FDL tendon (pink) and scar tissue (blue) at day 14 post-surgery. (E) 3D Reconstruction of segmented tissues at day 14 post-surgery. (E’) 3D reconstruction and volumetric quantification of STV at day 14 post-surgery.

*Segmentation and validation with histology:* B-mode images were exported as 3D volumes and loaded in to AMIRA (FEI v.6.1.1, Thermo Scientific, Hillsboro, OR). The scar tissue boundaries were identified and segmented on each slice. A 3D reconstruction of the scar tissue was generated and volumetrically quantified, resulting in the Scar Tissue Volume (STV) metric. To validate the correct segmentation of STV in US images, a subset of specimens (n=5) underwent both US imaging and histological evaluation. Following US imaging, mice were sacrificed, and hind paws were harvested and fixed in 10% neutral buffered formalin (NBF) for 72 hours at room temperature. Samples were then decalcified for 7 days in 14% EDTA^15^ at room temperature and processed for paraffin histology. Serial 5am transverse sections were cut through the entire width of the hindpaw that included the flexor tendon and/or scar tissue. For 3D reconstruction, sections corresponding to every 40em were stained with Alcian Blue/ Hematoxylin/ Orange G (ABHOG), as the step size of the US images was 40im. ABHOG was used as it allows easy discrimination between native tendon and scar tissue^16^. Stained sections were then digitally imaged, aligned, and stacked using NIH Image J ^17^, and loaded in to AMIRA. Scar tissue was then manually segmented in each slice, and volumetrically quantified.

*Assessment of Gliding Function and Mechanical Testing*: Following ultrasound imaging, C57Bl/6J mice were sacrificed at day 14 or day 28 post-surgery (n=7 per time-point) for assessment of gliding function and tensile mechanical testing ^4,7,16^. The hindlimb was disarticulated at the knee and the skin was removed down to the ankle. The FDL tendon was isolated at the myotendinous junction and secured between two pieces of tape. The tibia was gripped in an alligator clip and the FDL was incrementally loaded with small weights from 0-19g. Digital images were taken after each weight was applied and the flexion angle at the metatarsophalangeal (MTP) joint was measured from these images. The MTP Flexion angle corresponds to the difference in flexion from the neutral, unloaded (0g) image, and the flexion angle when the 19g weight is applied. Application of a 19g weight results in complete flexion of uninjured FDL tendons. Gliding Resistance was calculated based on the changes in MTP Flexion Angle over the range of applied loads with higher Gliding Resistance indicating impaired gliding function. Following gliding assessment, tendons were released from the tarsal tunnel, the tibia and calcaneus were removed, and the repaired tendon underwent tensile testing as previously described ^4,7^. Briefly, the toes and the proximal end of the tendon were secured in opposing custom grips in an Instron 8841 uniaxial testing system (Instron Corporation, Norwood, MA) and tested in tension at a rate of 30mm/min, until failure.

*Statistical Analyses*: To identify significant differences in STV over time in C57BL/6J mice, a one-way Analysis of Variance (ANOVA) with post-hoc multiple comparisons was used. Student t-tests were used to identify significant differences between WT and S10a04^GFP/+^repairs. Significance was set at p<0.05. Two independent blinded observers performed all subjective readings (*e.g.* segmentation of STV, assessment of gliding function). Univariate regression analysis was used to determine if STV measured from ultrasound images correlated with gliding function or tensile mechanical properties. To evaluate intraoperator and interoperator error in the segmentation of STV, two operators measured 10 randomly selected C57BL/6J specimens. The average percent error was calculated as the absolute difference between measures divided by the average measurement. The coefficient of variance was calculated as previously described^18^. Percent error was also calculated between STV quantified from segmentation of histology and ultrasound in the same specimens.

## Results

### Segmentation and quantification of Scar Tissue Volume from ultrasound images

To determine whether tendon healing could be measured non-invasively using ultrasound, C57BL/6J mice underwent complete transection and repair of the Flexor Digitorum Longus (FDL) tendon, and ultrasound imaging was performed at 7, 14, 20, and 28 days post-surgery. High-frequency ultrasound permits the identification of the healing (Figure 1C) FDL tendon in 2D sagittal B mode images. From these images, discreet tissues can be segmented on each 2D slice (Figure 1D), including native tendon (pink), bone (green), skin (yellow) and scar tissue (blue). Following segmentation, tissues were reconstructed in 3D (Figure 1E), and Scar Tissue Volume (STV) was quantified (Figure 1E’). To determine the potential intra- and interoperative sources of error in the quantification of STV. The average percent error between operators’ measurements was 15.6%, while the coefficient of variation was 16.6%.

### Histological validation of US segmentation of STV

To confirm that US image segmentation for volumetric measurement was accurately identifying scar tissue we analyzed 3D reconstructions of histological and US images from the same specimens and observed comparable morphology when segmenting images from both modalities. In addition, volumetric quantification of STV between modalities was highly consistent with a 10.8% error between US and histology segmentation. Taken together, these data suggest our ability to properly identify and segment scar tissue via US imaging (Fig. 2A-H).

**Figure 2.**
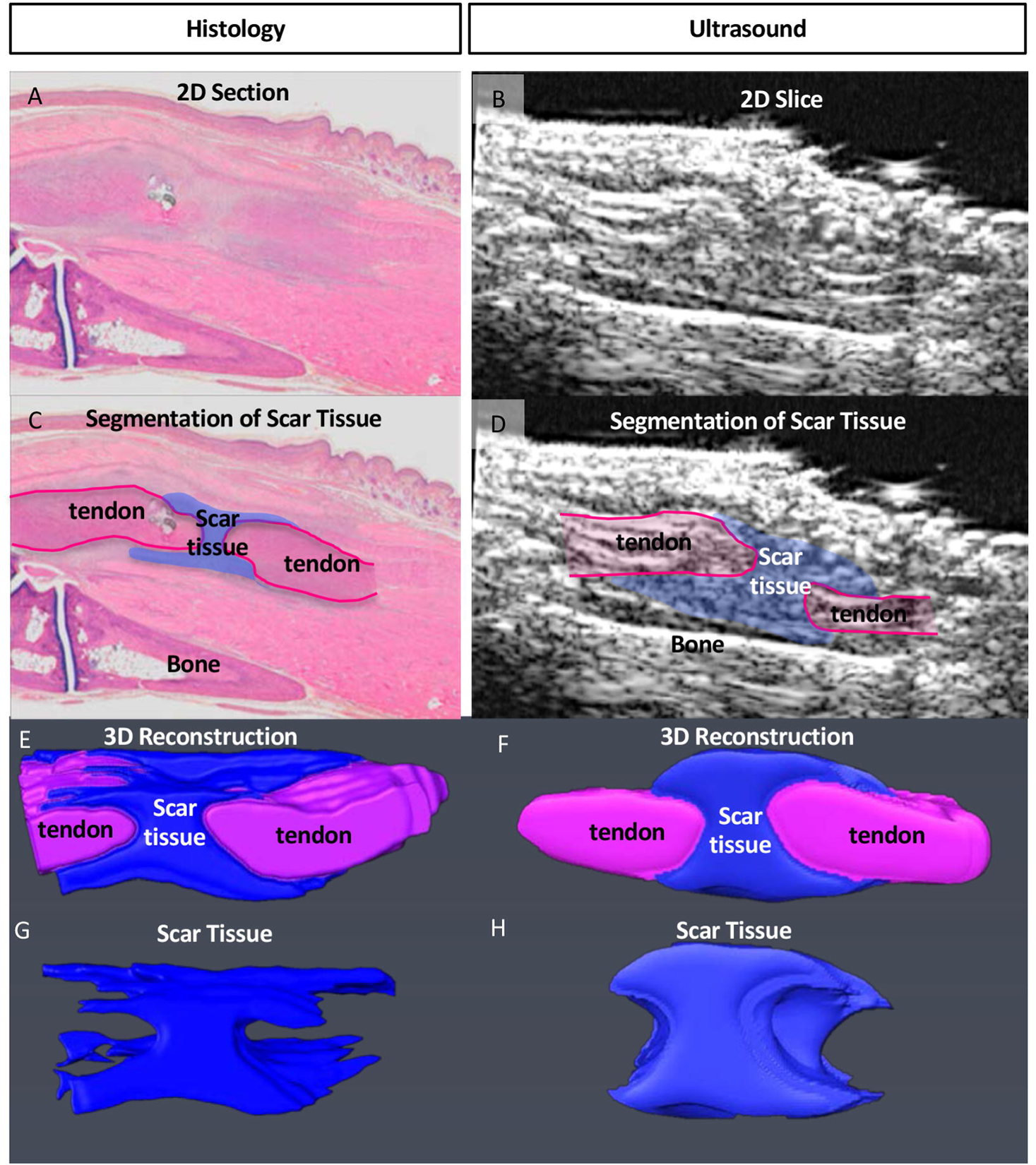
Validation of Scar Tissue Segmentation using histology. (A & B) 2-Dimensional transverse Histological (A) and US (B) image sections from the same specimen. (C & D) Segmentation of scar tissue (blue) and tendon (pink) from (C) histology, and (D) US images. (E & F) Following segmentation of all 2D images containing scar tissue from (E) histology, and (F) US, the segmented slices were reconstructed in 3D, and (G & H) scar tissue was volumetrically quantified.

### Scar Tissue Volume is strongly correlated with end-point metrics of gliding function

No detectable scar tissue was observed in un-injured tendons. At day 7 post-surgery average STV was 0.96 ± 0.158mm^3^, with a subsequent increase on day 14 (1.29 mm^3^ ± 0.10). Peak STV was observed at day 20 (1.55 mm^3^ ± 0.19) with a significant decrease observed at day 28 (0.91 mm^3^ ± 0.07), relative to day 20 (p<0.05) (Figure 3A). To determine the relationship between STV and metrics of gliding function, univariate linear regression analyses were performed. When day 14 and 28 data were grouped, a significant inverse correlation was observed between STV and MTP Flexion Angle (R^2^=0.070, p=0.0002) (Figure 3B), while a significant, positive correlation was observed between STV and Gliding Resistance (R^2^=0.63, p=0.0007) (Figure 3C). However, when each timepoint was analyzed separately, stronger correlations were observed between STV and gliding function at Day 14 and weaker, non-significant correlations were observed at day 28.

**Figure 3.**
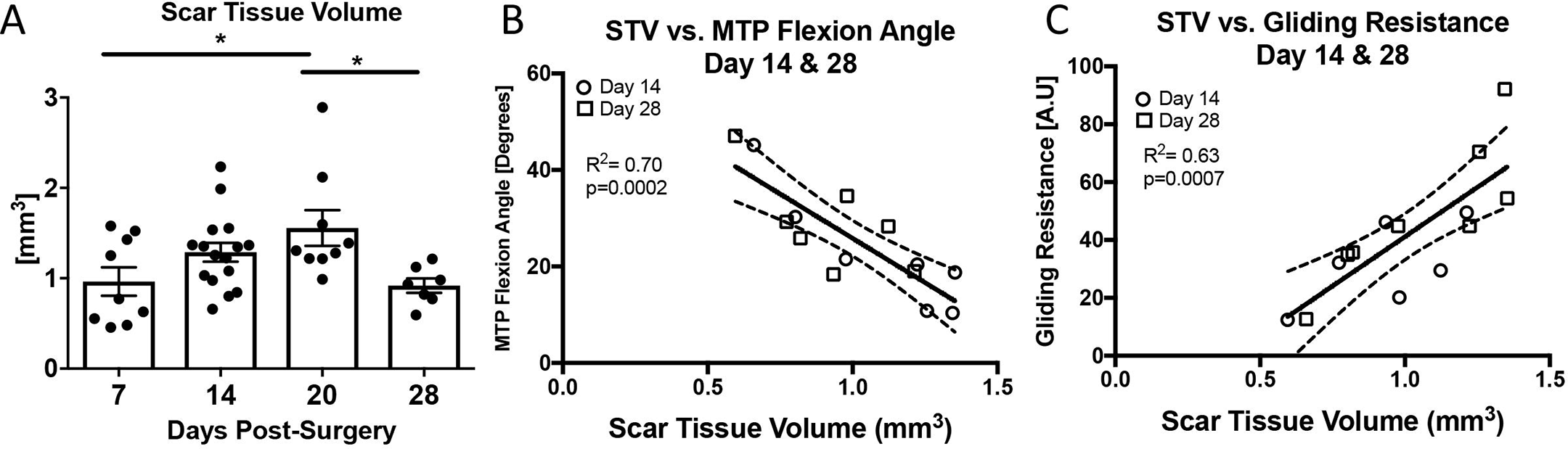
Scar Tissue Volume is correlated with changes in gliding function. (A) Scar Tissue Volume was quantified longitudinally at 7, 14, 20, and 28 days post-surgery. Peak STV was observed at day 20. (*) indicates p<0.05. (B-C) Linear regression analyses of STV and (B) MTP Flexion Angle, (C) Gliding Resistance at 14 and 28 days. Circles represent Day 14 and squares represent day 28.

At Day 14, a strong, significant inverse correlation was observed between STV and MTP Flexion Angle (R^2^=0.82, p=0.005), while a strong, significant positive correlation was observed between STV and Gliding Resistance (R^2^=0.71, p=0.01). At Day 28 weak, non-significant correlations were observed between STV and MTP Flexion Angle (R^2^=0.46, p=0.09) and Gliding Resistance (R^2^=0.36, p=0.15) (Table 1). Taken together, these data suggest that STV is a significant indicator of gliding function during the earlier phases of tendon healing, but not during later healing.

**Table 1.** Correlations between Scar Tissue Volume and Functional Metrics at 14 and 28 days post-surgery.

|  | Scar Tissue Volume |  |  |  |
| --- | --- | --- | --- | --- |
|  | Day 14 |  | Day 28 |  |
|  | r | p value | r | p value |
| Gliding/ Biomechanics parameter |  |  |  |  |
| Gliding Resistance | 0.71** | 0.01 | 0.36 | 0.15 |
| MTP Flexion Angle | 0.82** | 0.005 | 0.46 | 0.09 |
| Stiffness | 0.54 | 0.06 | 0.009 | 0.83 |
| Max load at failure | 0.09 | 0.49 | 0.25 | 0.24 |

### Scar Tissue Volume does not predict tensile mechanical properties

To determine the potential relationship between STV and tensile mechanical properties, univariate linear regression analyses of STV and tensile mechanical properties (Stiffness, Max load at failure) were conducted. When days 14 and 28 were grouped there were no significant correlations between STV and Stiffness (R^2^=0.13, p=0.19) (Figure 4A) or Max load (R^2^=0.05, p=0.68) (Figure 4B). Furthermore, no significant associations were observed when each time-point was analyzed separately (Table 1), although a moderately strong but non-significant association was observed between STV and Stiffness at D14 (R^2^=0.54, p=0.06).

**Figure 4.**
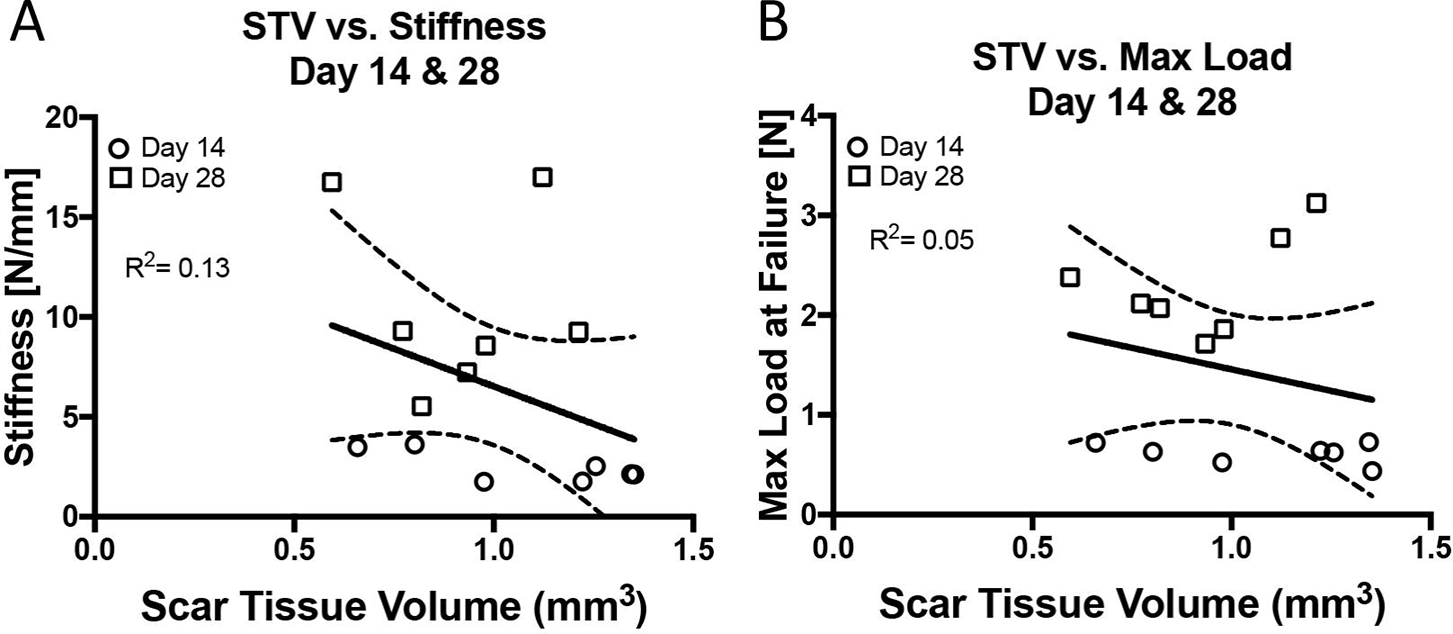
Scar Tissue Volume is not correlated with tensile mechanical properties. Linear regression analyses of STV and (A) Stiffness, (B) Max load at failure at 14, and 28 days post-surgery. Circles represent Day 14 and squares represent day 28.

### STV identifies differences in models of fibrotic vs. regenerative healing

We have recently demonstrated that S100a4 haploinsufficiency (S100a4^GFP/+^) results in improved gliding function and mechanical properties, relative to wildtype (WT) littermates^19^. To demonstrate the potential of STV to identify regenerative versus fibrotic models of tendon healing, STV was quantified at day 14 post-surgery. A significant 27% decrease in STV was observed in S100a4^GFP/+^tendons, relative to WT (p=0.0053) (Figure 5), suggesting that STV has the sensitivity to non-invasively identify functional differences between modes of healing.

**Figure 5.**
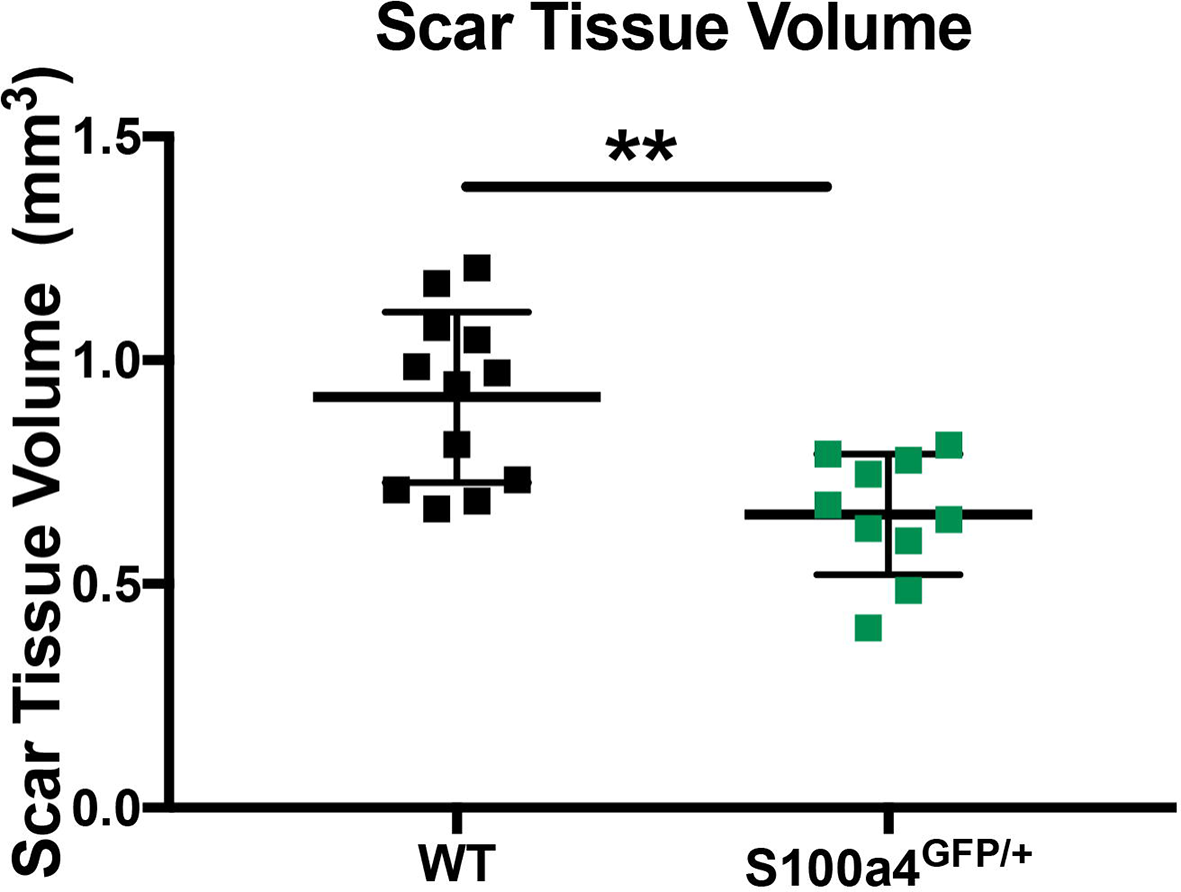
Scar Tissue Volume identifies functional differences between models of scar-mediated and regenerative tendon healing. Quantification of Scar Tissue Volume in WT and S1004 haploinsufficient (S100a4^GFP/+^) tendon repairs at day 14 post-surgery. S100a4^GFP/+^mice heal with improved gliding function and decreased STV. (**) indicates p<0.001.

## Discussion

Following injury, tendons are prone to a fibrotic, scar mediated healing process that both impairs restoration of range of motion and hinders the reacquisition of normal mechanical properties. Given that there are currently no biological approaches to improve the tendon healing process, there is a massive need for increased pre-clinical screening and identification of potential therapies. To address, we have identified Scar Tissue Volume as a non-invasive ultrasound metric to determine the effects of genetic or pharmacological modifications on the healing process, which may allow more rapid identification of tendon therapeutics. Importantly, this approach has the potential to dramatically decrease the number of animals needed, consistent with the goals of reducing the number of animals used in biomedical research^20^. Moreover, non-invasive, longitudinal ultrasound imaging can promote more rapid, cost-effective and high-throughput screening to facilitate the identification of therapeutic targets.

Several studies have previously demonstrated the potential of using ultrasound to non-invasively image the healing tendon. Ghorayeb *et al*., quantified extracellular matrix content and found a strong correlation between ECM content and linear stiffness in the Achilles tendon, although changes in range of motion were not assessed. In the current study, we did not observe a significant association between STV and tissue stiffness, although there was a moderately strong relationship at day 14 (R^2^=0.54). While quantification of ECM and STV may be similar in that both metrics are likely related to tissue size or bulk, which are related to impairments in range of motion, differences in correlation with stiffness between ECM and STV may be due to differences in the tissues that were quantified for ECM and STV, timing of analysis, or models of healing. In contrast to quantification of tissue content or volume, several studies have examined the relationship between ultrasound echogenicity and healing. Tamura *et al*., used US echogenicity to assess healing in equine Superficial Digital Flexor Tendons, although no change in echogenicity was observed despite longitudinal improvements in strain. More recently, Lee *et al*., assessed the relationship between US echogenicity, and tensile mechanical properties in a collagenase-induced tendinopathy model and demonstrated that echo intensity was positively correlated with maximum strain and stiffness^21^. Interestingly, tendon cross-sectional area and echogenicity were increased in healing tendons, relative to controls. Although no correlation analysis was conducted between these parameters, future studies will be needed to understand the potential relationship between STV and echo intensity.

An important aspect of this approach is the ability to detect functional differences related to tendon ROM non-invasively and over time. However, gait analysis can also be used to non-invasively assess restoration of function during tendon healing^22-24^. While gait analysis is a strong metric in assessing Achilles and supraspinatus tendon healing in pre-clinical models, it is not yet known how alterations in gait correspond to flexor tendon healing phases. Moreover, gait analysis parameters are typically associated with measurements of pain and weakness, while the relationship between gait and tissue morphology are unknown. While we observe changes in STV over the course or healing, these differences are relatively modest, with no differences in STV observed between 14- and 20-days post-surgery. However, we typically observe only slight differences in gliding function metrics between these time-points^4^. Importantly, the real utility of end-point metrics of gliding function is in identifying differences between genetically different strains of mice^10,16^, or between pharmacological treatment groups^11^ at a given timepoint. Taken together, these data strongly suggest that longitudinal US based quantification of scar tissue volume may be utilized as a corollary to predict tendon ROM and gliding function during healing.

While STV strongly correlates with gliding function, there are several limitations to this study that must be considered. STV does not correlate with mechanical properties, which are a critical indicator of the value of a particular therapeutic intervention. Thus, future work seeks to develop multiple US-based metrics which either alone or using multi-variate analyses may better correlate with restoration of mechanical properties. Furthermore, we have only assessed healing in S100a4^GFP/+^and WT mice at Day 14. However, we have confined our analysis in these animals to day 14 due to the decrease in predictive power of STV at day 28. Finally, while the goal of developing longitudinal, non-invasive metrics of healing is to permit rapid and cost-effective therapeutic target screening, the segmentation process requires substantial expertise and is quite time-consuming, thus to make this approach more high-throughput we will need to semi-automate or automate the segmentation process.

Here we have shown that Scar Tissue Volume is significantly correlated with end-point metrics of gliding function, and STV is particularly predictive of gliding function at Day 14, the period of peak impairments in gliding function. In addition, STV is sensitive enough to discriminate between phenotypically distinct models of healing. Future work will develop additional US-based metrics with the goal of non-invasive estimation of mechanical properties. Taken together, this study identifies quantification of Scar Tissue Volume using longitudinal, non-invasive ultrasound imaging as a novel means to assess tendon healing. This approach may permit the rapid screening of biological and pharmacological interventions for healing, and identify promising therapeutic targets, in an efficient, cost-effective manner.

